# EPHX1 Methylation, Tobacco Exposure, and Survival in Head and Neck Squamous Cell Carcinoma

**DOI:** 10.64898/2026.11.25.769160

**Authors:** Carlos Luis Pérez Hernández, Alejandro Rivero Bandinez

**Affiliations:** Centro de Desarrollo de las Ciencias Básicas Biomédicas, Universidad de Ciencias Médicas de La Habana (UCMH); Department of Biochemistry. Instituto de Ciencias Básicas y Preclínicas "Victoria de Girón" (ICBP), Universidad de Ciencias Médicas de La Habana (UCMH); Centro Estatal para el control de los Medicamentos y Equipos Médicos, CECMED

**Keywords:** HNSC, EPHX1, DNA methylation, Cancer, epigenetics, gene expression

## Abstract

**Background:** Epoxide hydrolase 1 (EPHX1) metabolizes tobacco-derived and environmental epoxides and has been implicated in carcinogen activation/detoxification. Whether EPHX1 methylation is associated with cigarette smoking, tumor status, gene expression, and survival in head and neck squamous cell carcinoma (HNSC) remains incompletely defined.

**Methods:** We analyzed TCGA-HNSC DNA methylation data for EPHX1 CpGs, together with smoking exposure (years smoked), EPHX1 expression, tumor/normal status, and clinical follow-up. Methylation was measured as M-values and Beta-values. Associations between smoking duration and CpG methylation were tested using linear regression. Tumor-normal differential methylation and expression were assessed using linear models. Methylation-expression correlations were evaluated using Pearson correlation. Kaplan-Meier analyses and log-rank tests were used to examine overall survival (OS).

**Results:** Longer smoking duration was significantly associated with decreased methylation (hypomethylation) at five EPHX1 CpGs in the tumor cohort, most notably at cg26513734 (P = 0.0018) and cg05935800 (P = 0.0118). No significant associations were observed in normal tissue. Four EPHX1 CpGs demonstrated a significant negative correlation with EPHX1 gene expression (r = −0.556 to −0.372, all P < 0.001). EPHX1 was differentially methylated between tumor and normal tissue, most notably at cg03138928 (P = 1.78e-32). High methylation at cg05935800 (P = 0.0016), cg02453929 (P = 0.044), and cg24868305 (P = 0.048) was associated with significantly worse overall survival.

**Conclusions:** These findings suggest that tobacco exposure is associated with tumor-specific EPHX1 hypomethylation, which in turn is linked to altered gene expression and poorer clinical outcomes.

## Introduction

Head and neck squamous cell carcinoma (HNSC) represents a major global health burden. In 2022, 947,211 new cases and 482,428 deaths were reported worldwide, accounting for 4.7% of all cancer diagnoses and 4.9% of all cancer-related mortalities. If the trend continues, the incidence of HNSC is projected to rise substantially in the coming decades, with estimates suggesting an increase to 1,550,796 new cases and 823,471 deaths in 2050, representing an increase of 1.6 and 1.7 times, respectively. The disease encompasses malignancies arising from the oral cavity, pharynx, and larynx, and is characterised by significant heterogeneity in terms of aetiology, molecular pathogenesis, and clinical outcomes. (1–3)

The principal risk factors for HNSC are well established and include tobacco use, alcohol consumption, and infection with high-risk human papillomavirus (HPV). Tobacco smoking and alcohol drinking are major risk factors for HNSC, accounting for up to 72% of cases, with slight differences between subtypes. Tobacco smoking poses the greatest risk for laryngeal cancer, whereas alcohol drinking poses the greatest risk for cancer of the oral cavity and oropharynx, and high-risk HPV is a major risk factor for oropharyngeal cancer. (4,5)

Tobacco smoke contains a complex mixture of procarcinogens, including polycyclic aromatic hydrocarbons (PAHs) and aromatic amines, which require metabolic activation to exert their mutagenic effects. Microsomal epoxide hydrolase 1 (EPHX1), encoded by the EPHX1 gene, is a critical phase I biotransformation enzyme that catalyses the hydrolysis of epoxide derivatives generated during this metabolic processing. EPHX1 plays a dual role in carcinogenesis, functioning in both the detoxification and, in some contexts, the activation of reactive epoxide intermediates. EPHX1 genetic polymorphisms and modified expression have been studied in relation to cancer susceptibility, including HNSC. (6–10)

Beyond genetic variation, epigenetic mechanisms—particularly DNA methylation— provide a major regulatory layer that controls gene expression, including EPHX1 and, by extension, cellular detoxification capacity. Promoter CpG island hypermethylation is a canonical mechanism of transcriptional silencing in cancer, and its role in inactivating tumour suppressor and DNA repair genes is well documented. In the context of EPHX1, evidence shows that EPHX1 methylation has been linked to its expression in several conditions, including cancer, supporting the functional relevance of this epigenetic mark.(8,11–13) However, the specific landscape of EPHX1 CpG methylation and its relationship to tobacco exposure in HNSC has not been comprehensively defined.

The clinical relevance of DNA methylation as a prognostic biomarker in head and neck cancer is increasingly recognised. A recent systematic review and meta-analysis confirmed that high levels of DNA methylation are consistently associated with poor overall survival and disease-free survival in head and neck cancer, providing robust evidence for the utility of methylation-based markers. (14,15) Despite these advances, locus-specific studies focusing on genes with established roles in carcinogen metabolism, such as EPHX1, remain sparse. Elucidating how tobacco exposure shapes the EPHX1 methylation profile, whether this methylation differs between tumour and normal tissue, and whether it influences gene expression and patient survival could provide mechanistic insight into the pathogenesis of tobacco-associated HNSC.

The present study was therefore designed to address these gaps by integrating multi-omic data from The Cancer Genome Atlas (TCGA) HNSC cohort. We systematically examined the associations between smoking duration and EPHX1 CpG methylation, characterised tumour-normal differential methylation across the locus, evaluated the correlation between methylation and EPHX1 mRNA expression, and assessed the impact of EPHX1 methylation on overall survival. We hypothesised that tobacco exposure would be associated with altered EPHX1 methylation, that these epigenetic changes would be tumour-specific and inversely correlated with gene expression, and that high methylation at prognostic CpG sites would confer an increased risk of death.

## Materials and Methods

### Data Source and Cohort

Publicly available multi-omic data for Head and Neck Squamous Cell Carcinoma (TCGA-HNSC) were obtained from the Genomic Data Commons (https://www.cancer.gov/tcga.). The dataset included Illumina 450K DNA methylation arrays, RNA-sequencing data, and matched clinical annotations (including survival and self-reported "years smoked"). Samples were analysed as primary solid tumours and solid tissue normals. Sample sizes varied across analyses based on data completeness.

### Methylation Data Processing

Methylation was quantified using Beta-values (β, 0–1) and M-values (logit-transformed β). M-values were used for all statistical modelling due to their homoscedasticity, which reduces false positives at extreme methylation levels, while Beta-values were retained for biological interpretation and graphical display. (16)

### Smoking Association Analysis

Smoking exposure was defined as cumulative "years smoked," a continuous variable that retains statistical power compared to categorical smoking status. Linear regression models were used to test the association between years smoked and CpG M-values, independently in tumour and normal tissues. Linear regression was chosen to estimate the effect size (change in methylation per year) and identify tumour-specific exposure effects.

### Differential Methylation Analysis

Tumour-normal differential methylation was assessed using linear modelling (via the limma package) of M-values, providing moderated t-statistics to stabilise variance across CpGs. Effect sizes are reported as Delta M (ΔM): positive values indicate tumour hypermethylation, and negative values indicate tumour hypomethylation.

### Methylation–Expression Correlation

Pearson correlation coefficients (r) were calculated between CpG Beta-values and log₂-transformed EPHX1 mRNA expression to assess the functional impact of methylation. Pearson correlation was selected to quantify the linear direction and strength of the relationship, with negative r values indicating methylation-mediated transcriptional silencing.

### Survival Analysis

Overall survival (OS) was defined as time from diagnosis to death from any cause, with living patients censored at last follow-up. Kaplan–Meier curves, stratified by median methylation (high vs. low), were compared using the non-parametric log-rank test to assess survival differences without distributional assumptions. Univariate Cox proportional hazards regression was used to estimate Hazard Ratios (HRs) and 95% confidence intervals (CIs), where HR > 1 indicates an increased risk of death.

### Statistical Correction and Software

To account for multiple testing, the Benjamini–Hochberg false discovery rate (FDR) procedure was applied, controlling the expected proportion of false positives while preserving statistical power for biologically relevant findings. (17) All analyses were performed in R using the limma, survival, and survminer packages.

### Ethical Considerations

This study used publicly available, de-identified data from The Cancer Genome Atlas (TCGA) Head and Neck Squamous Cell Carcinoma (TCGA-HNSC) project, accessed through the Genomic Data Commons. Because the dataset is fully anonymised and publicly accessible, this secondary analysis did not require additional institutional review board (IRB) approval or informed consent. The original TCGA study was conducted in accordance with the Declaration of Helsinki and was approved by the ethics committees of all participating institutions.

## Results

### Summary of EPHX1 CpG Alterations

Table 1 provides a comprehensive summary of the differential methylation (Tumour vs. Normal), survival associations, and smoking associations for all analysed CpGs, ordered by tumour-normal differential methylation significance.

**Table 1.** Summary of EPHX1 CpG methylation alterations across tumour-normal comparisons, overall survival, and smoking exposure in Head and Neck Squamous Cell Carcinoma (HNSC).

| **CpG Probe** | **Delta M (T vs N)** | **P (T vs N)** | **P (Survival)** | **P (Smoking)** |
|---|---|---|---|---|
| cg03138928 | 1.633 | 1.78e-32 | 8.37e-01 | 5.49e-01 |
| cg05935800 | 0.909 | 3.80e-11 | 1.59e-03 | 1.18e-02 |
| cg02453929 | 0.299 | 2.24e-05 | 4.44e-02 | 4.73e-01 |
| cg03459809 | 0.455 | 2.68e-04 | 4.26e-01 | 4.44e-01 |
| cg26513734 | -0.581 | 5.48e-04 | 2.36e-01 | 1.80e-03 |
| cg24868305 | 0.466 | 1.15e-03 | 4.75e-02 | 8.40e-02 |
| cg05385434 | 0.562 | 2.03e-03 | 1.26e-01 | 2.22e-02 |
| cg11135003 | 0.093 | 1.13e-02 | 1.70e-01 | 5.52e-01 |
| cg15633273 | -0.434 | 1.31e-02 | 6.77e-01 | 1.89e-02 |
| cg24928687 | -0.233 | 1.40e-02 | 3.74e-01 | 2.88e-01 |
| cg00501904 | 0.510 | 1.68e-02 | 5.03e-01 | 2.25e-02 |
| cg23096144 | 0.186 | 1.95e-02 | 2.78e-01 | 9.46e-01 |
| cg25152404 | 0.338 | 2.64e-02 | 9.35e-01 | 9.39e-02 |
| cg00396625 | 0.070 | 4.52e-02 | 9.66e-01 | 5.64e-01 |
| cg25004581 | 0.178 | 8.22e-02 | 3.15e-01 | 6.37e-01 |
| cg26187962 | -0.162 | 8.47e-02 | 5.91e-01 | 3.23e-01 |
| cg23992403 | -0.174 | 9.48e-02 | 1.56e-01 | 2.58e-01 |
| cg17468616 | 0.123 | 1.63e-01 | 3.97e-01 | 6.97e-01 |
| cg20316779 | -0.059 | 2.00e-01 | 7.04e-01 | 7.96e-01 |
| cg11967292 | 0.040 | 3.65e-01 | 6.52e-01 | 9.88e-01 |
| cg17442483 | -0.148 | 3.98e-01 | 9.28e-01 | 1.57e-01 |
| cg24350360 | 0.037 | 6.21e-01 | 7.40e-01 | 1.30e-01 |
| cg03337430 | -0.022 | 7.48e-01 | 3.31e-01 | 9.14e-01 |
| cg05605921 | 0.014 | 7.72e-01 | 6.11e-01 | 7.24e-01 |

Table 1 presents the statistical summary for all analysed EPHX1 CpG probes in the TCGA-HNSC cohort, ordered by the significance of the tumour-normal differential methylation (P (T vs N)). Statistical significance for all tests was defined as a two-sided P-value < 0.05. CpG Probe: Illumina identifier for the specific cytosine-phosphate-guanine site. Delta M (T vs N): Difference in mean methylation M-values between primary solid tumour samples and solid tissue normal samples (Tumour minus Normal). Positive values indicate hypermethylation in tumours relative to normal tissue; negative values indicate hypomethylation in tumours. P (T vs N): P-value from a linear model assessing differential methylation between tumour and normal tissue. P (Survival): P-value from a log-rank test comparing overall survival (OS) between patients stratified by high (above median) versus low (below median) methylation at the respective CpG. P (Smoking): P-value from a linear regression model testing the association between cumulative years smoked and methylation M-values within the tumour cohort. Abbreviations: M-value, logit-transformed beta-value; T vs N, Tumour vs Normal.

### Longer smoking duration is associated with EPHX1 hypomethylation specifically in tumour tissue

We evaluated the relationship between smoking duration (years smoked) and *EPHX1* methylation. In the tumor cohort, longer smoking duration was significantly associated with **decreased methylation (hypomethylation)** at five distinct CpGs: cg26513734 (Estimate = −0.017, P = 0.0018), cg05935800 (Estimate = −0.014, P = 0.0118), cg15633273 (Estimate = −0.014, P = 0.0189), cg05385434 (Estimate = −0.017, P = 0.0222), and cg00501904 (Estimate = −0.020, P = 0.0225). Notably, this association was absent in the normal tissue cohort (all P > 0.05), indicating that tobacco-induced epigenetic alterations at the *EPHX1* locus are tumour-specific (Figure 1).

**Figure 1.**
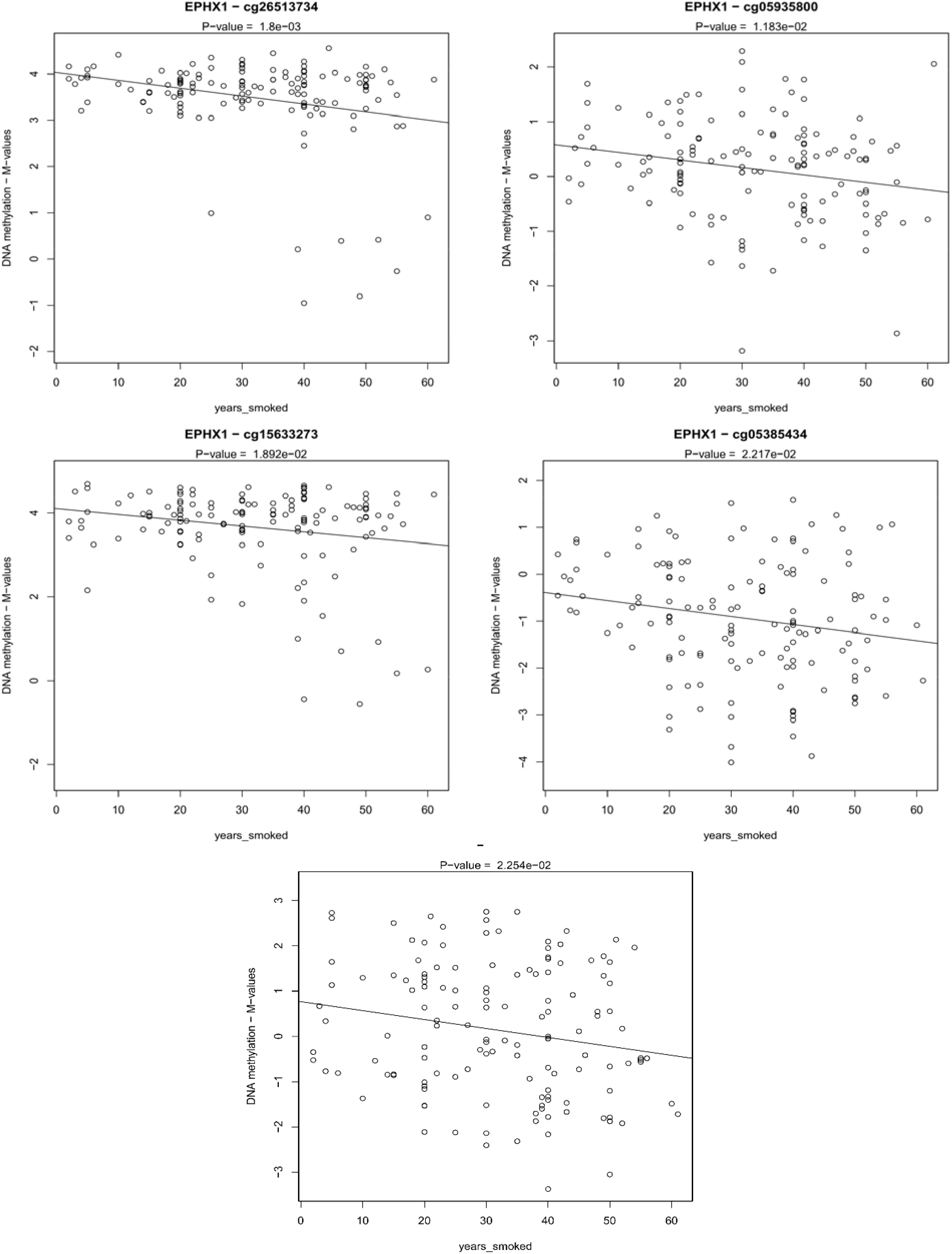
Association between cumulative smoking duration and EPHX1 CpG methylation in Head and Neck Squamous Cell Carcinoma (HNSC) tumour tissue. Scatter plots display the relationship between self-reported cumulative years smoked (x-axis) and DNA methylation levels, expressed as M-values (y-axis), for five EPHX1 CpG sites in primary solid tumour samples from the TCGA-HNSC cohort. Each open circle represents an individual patient sample. Solid black lines denote the linear regression best-fit lines. A negative slope indicates an inverse relationship. P-values displayed above each plot were derived from linear regression models testing the association between years smoked and methylation; a two-sided P-value < 0.05 was considered statistically significant. Abbreviations: M-values, logit-transformed beta-values (a continuous metric where negative values indicate lower methylation and positive values indicate higher methylation).

### EPHX1 is significantly differentially methylated between tumour and normal tissue

Analysis of primary solid tumours versus solid normal tissue revealed widespread differential methylation. Notably, we observed profound **hypermethylation in tumours** at multiple loci, most significantly at cg03138928 (Delta M = +1.633, P = 1.78e-32), cg05935800 (Delta M = +0.909, P = 3.80e-11), cg03459809 (Delta M = +0.455, P = 2.68e-4), cg02453929 (Delta M = +0.299, P = 2.24e-5), and cg05385434 (Delta M = +0.562, P = 2.03e-3). Conversely, we identified significant **hypomethylation in tumours** at cg26513734 (Delta M = −0.581, P = 5.48e-4) and cg15633273 (Delta M = - 0.434, P = 1.31e-2) (Figure 2).

**Figure 2.**
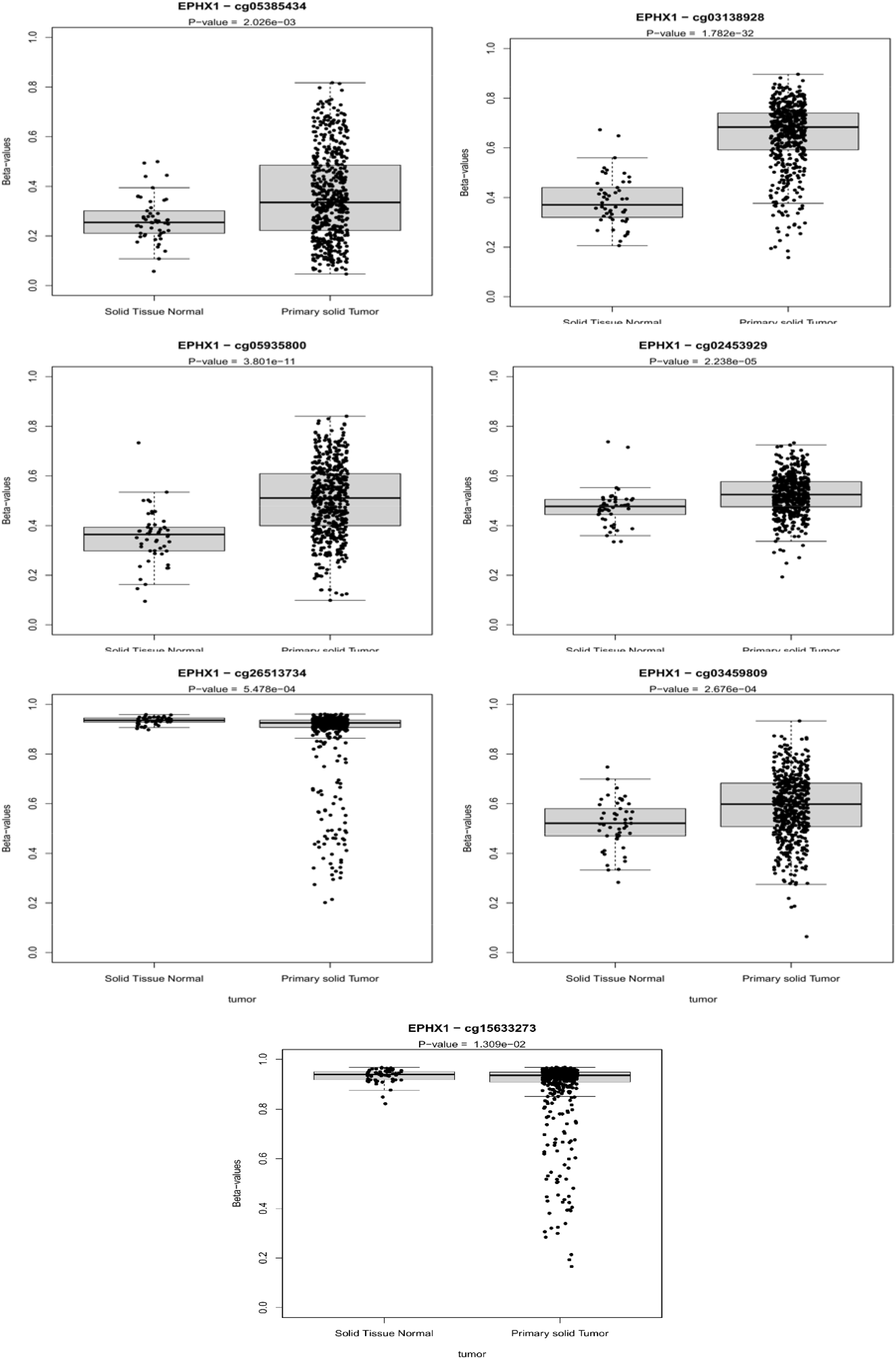
Differential DNA methylation of EPHX1 CpG sites between primary solid tumours and solid tissue normals in Head and Neck Squamous Cell Carcinoma (HNSC). Boxplots illustrate the distribution of DNA methylation levels at seven distinct EPHX1 CpG sites in adjacent solid tissue normal samples (left group) compared to primary solid tumour samples (right group) from the TCGA-HNSC cohort. Methylation is displayed as Beta-values on the y-axis, which represent the ratio of methylated probe signal to total signal, ranging from 0 (completely unmethylated) to 1 (completely methylated). In each boxplot, the solid horizontal line within the box represents the median methylation level, the box boundaries represent the interquartile range (IQR, 25th to 75th percentiles), and the whiskers extend to 1.5 times the IQR. Individual patient samples are overlaid as jittered black dots to display the full distribution of the data. P-values shown above each plot were calculated using linear regression models comparing methylation M-values (logit-transformed Beta-values) between the two tissue types. A two-sided P-value < 0.05 was considered statistically significant.

### EPHX1 methylation is strongly and negatively correlated with EPHX1 gene expression

To determine the functional impact of these epigenetic changes, we correlated methylation with *EPHX1* mRNA expression. Four CpGs demonstrated a highly significant **inverse correlation**, indicating that increased methylation leads to decreased gene expression: cg00501904 (r = −0.556, P < 0.001), cg03138928 (r = −0.526, P < 0.001), cg05385434 (r = −0.435, P < 0.001), and cg02453929 (r = −0.372, P < 0.001) (Figure 3).

**Figure 3.**
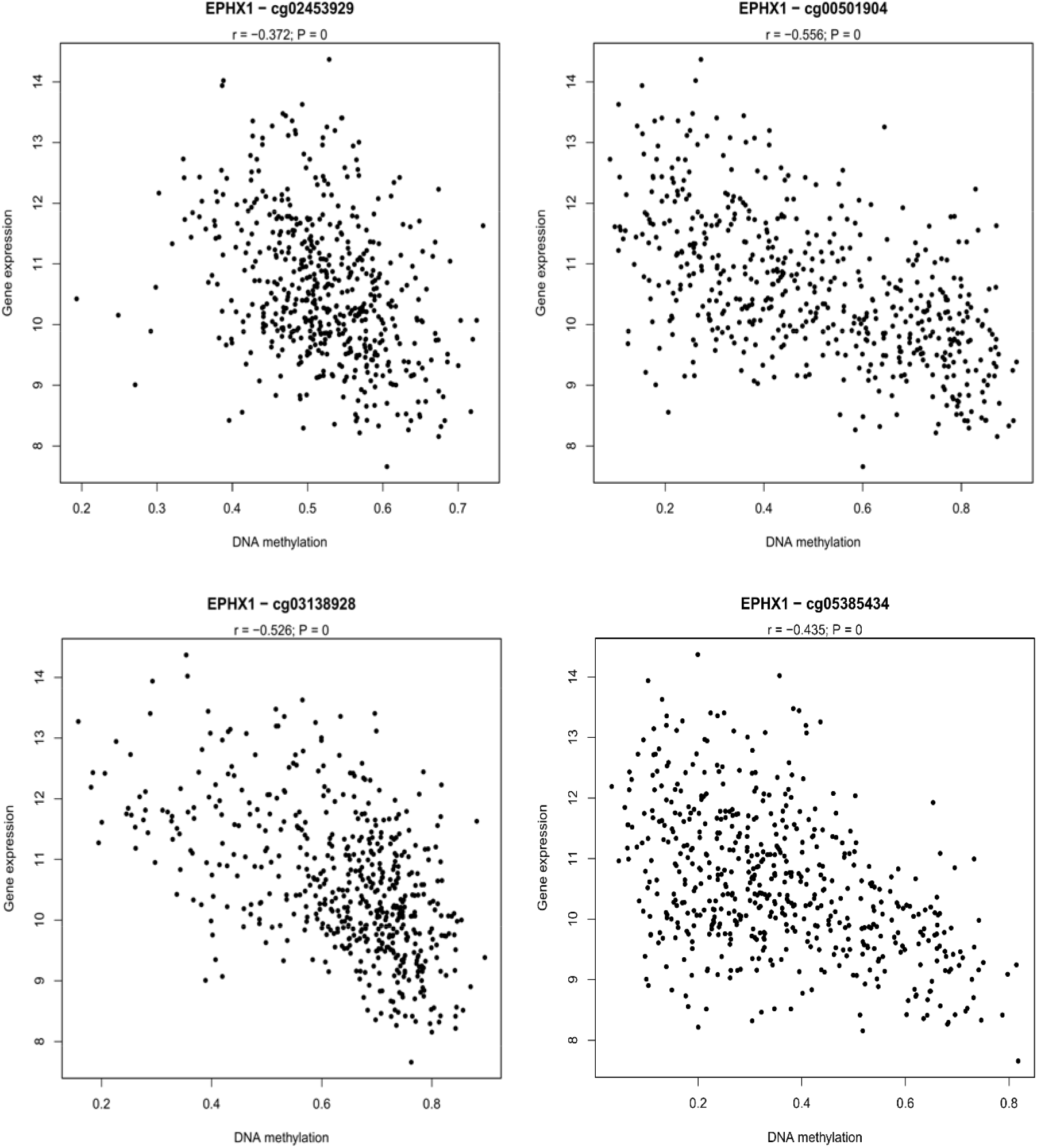
Correlation between EPHX1 CpG methylation and EPHX1 mRNA expression in Head and Neck Squamous Cell Carcinoma (HNSC). Scatter plots illustrate the relationship between DNA methylation at four individual EPHX1 CpG sites (x-axis) and EPHX1 gene expression (y-axis) in primary tumour samples from the TCGA-HNSC cohort. Each solid black dot represents an individual patient sample. DNA methylation is expressed as Beta-values, ranging from 0 (completely unmethylated) to 1 (completely methylated). Gene expression is displayed as log2-transformed mRNA levels. The strength and direction of the linear relationship were assessed using Pearson correlation analysis. The Pearson correlation coefficient (r) and associated two-sided P-value are displayed above each plot. Negative r values indicate an inverse correlation. A P-value < 0.05 was considered statistically significant (displayed as P = 0 due to numerical underflow, indicating P < 0.001).

### High EPHX1 methylation significantly increases the risk of poor overall survival

Kaplan-Meier survival analysis revealed that patients with **high methylation** at specific *EPHX1* loci face a significantly **increased risk of death** (worse overall survival) compared to those with low methylation. This was statistically significant for cg05935800 (Log-rank P = 0.0016), cg02453929 (Log-rank P = 0.044), and cg24868305 (Log-rank P = 0.048). The survival curves clearly demonstrate that the high-methylation groups (red lines) have steeper declines in survival over time (Figure 4). Other CpGs, including cg03138928 (P = 0.837) and cg05385434 (P = 0.126), did not show significant survival associations in this cohort.

**Figure 4.**
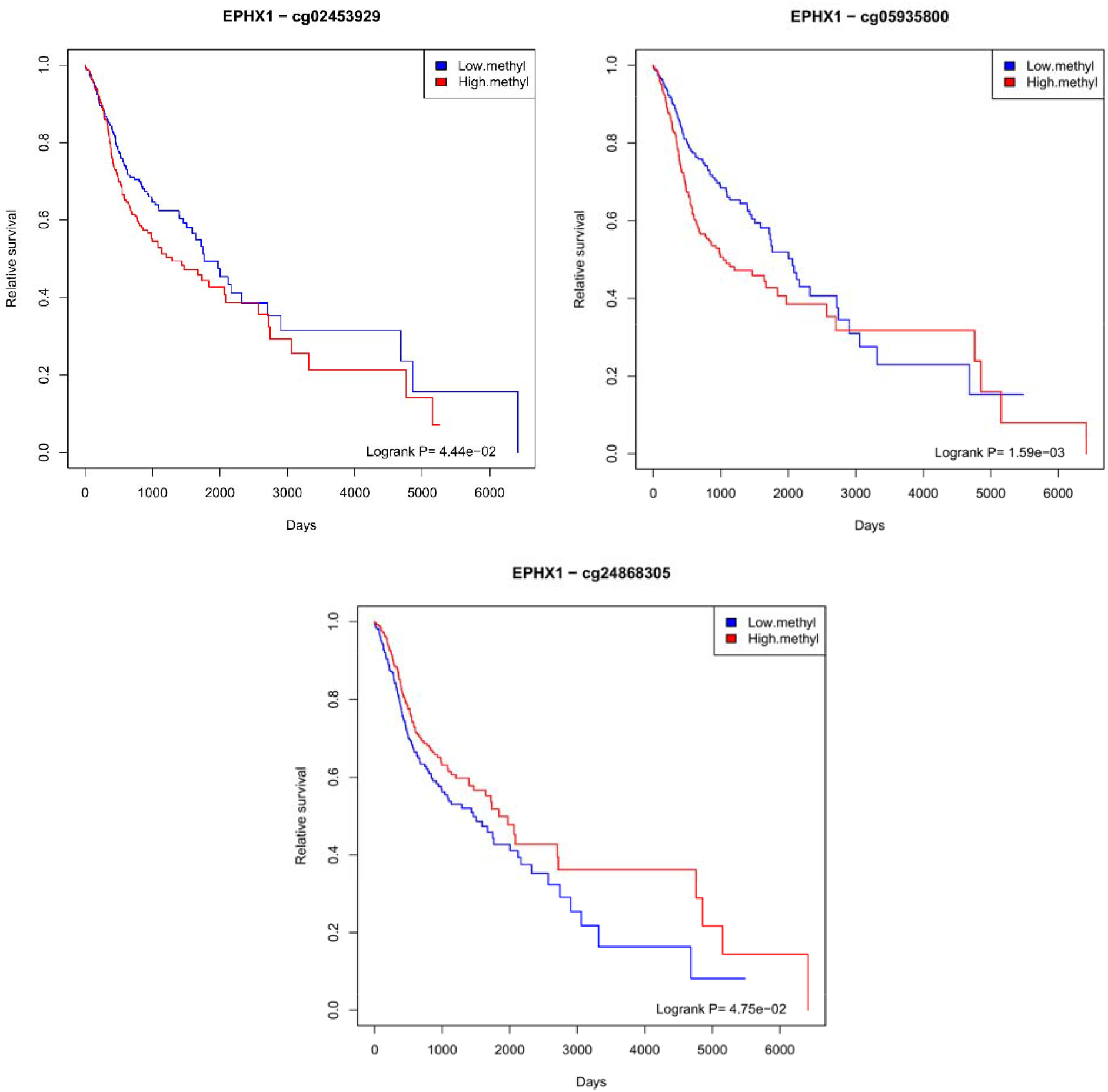
Kaplan-Meier overall survival analysis of Head and Neck Squamous Cell Carcinoma (HNSC) patients stratified by EPHX1 CpG methylation status. Kaplan-Meier curves display the probability of overall survival over time (x-axis, in days) for HNSC patients stratified into two groups based on EPHX1 DNA methylation levels at three specific CpG sites. Patients were categorised into "High methylation" (red lines) and "Low methylation" (blue lines) groups based on a median split of the methylation M-values. The y-axis represents the relative survival probability (ranging from 0.0 to 1.0). Vertical drops in the step functions indicate death events, while horizontal segments represent periods of survival; censored patients (those alive at last follow-up) are not explicitly marked with tick marks but are incorporated into the survival estimates. P-values displayed in the bottom right corner of each plot were calculated using the non-parametric log-rank test, which evaluates the null hypothesis that there is no difference in survival distributions between the high and low methylation groups. A two-sided P-value < 0.05 was considered statistically significant.

## Discussion

This integrative analysis of TCGA-HNSC data provides a multi-layered characterisation of EPHX1 epigenetic regulation and its clinical relevance in head and neck squamous cell carcinoma. The principal finding is that EPHX1 is subject to complex, context-dependent methylation control: smoking duration is associated with locus-specific hypomethylation in tumour tissue, tumours exhibit profound hypermethylation at discrete CpGs relative to normal tissue, and methylation at these loci is inversely correlated with EPHX1 expression and predictive of poor overall survival. Together, these observations position EPHX1 as a tobacco-responsive, methylation-regulated gene with prognostic significance in HNSC.

The functional rationale for studying EPHX1 in this context is well established. EPHX1 is a phase I biotransformation enzyme that catalyses the hydrolysis of epoxide derivatives of polycyclic aromatic hydrocarbons and aromatic amines generated from tobacco smoke, and it can act in either detoxification or metabolic activation depending on substrate and context.(18) Genetic variation in EPHX1 has been extensively investigated as a determinant of head and neck cancer susceptibility; a meta-analysis reported that the Tyr113His polymorphism may be a risk factor for HNC, while the His139Arg variant showed no association.(9) However, germline polymorphisms alone do not capture the dynamic, exposure-responsive regulation of gene expression that occurs during tumorigenesis. The present data suggest that epigenetic mechanisms— specifically CpG methylation—constitute an additional and potentially more plastic layer of EPHX1 regulation that is shaped by tobacco exposure and altered in the tumour microenvironment.

The observation that longer smoking duration is associated with decreased methylation at five CpGs in tumour tissue, but not in normal tissue, is particularly noteworthy. Prior studies in HNSCC have reported that smoking duration is associated with hypomethylation in tumours, suggesting that tobacco exposure leaves a distinct epigenetic imprint that may differ by etiologic subtype. (19,20) Our findings extend this paradigm to a carcinogen-metabolism gene, raising the possibility that smoking-induced hypomethylation of EPHX1 in tumour cells could alter detoxification capacity in a manner that influences tumour evolution. Importantly, the absence of this association in adjacent normal tissue argues against a simple constitutive effect of smoking on the EPHX1 locus and instead points to a tumour-specific interaction between exposure history and malignant transformation.

At the same time, the tumour-normal comparison revealed a contrasting pattern: profound hypermethylation at cg03138928, cg05935800, cg03459809, cg02453929, and cg05385434 in tumours relative to normal tissue, with cg03138928 exhibiting an exceptionally large effect (ΔM = +1.633, P = 1.78 × 10⁻³²). This dichotomy—smoking-associated hypomethylation at some loci and tumour-associated hypermethylation at others—suggests that the EPHX1 locus is subject to competing epigenetic pressures. One interpretation is that tumorigenesis drives widespread promoter hypermethylation as part of the canonical silencing of growth-regulatory and detoxification genes, while tobacco exposure may attenuate methylation at select CpGs. The convergence of these forces on cg05935800, which is both hypermethylated in tumours and hypomethylated with longer smoking duration, makes it a particularly compelling candidate for functional follow-up.

The functional consequence of these methylation changes is supported by the consistent inverse correlations between CpG methylation and EPHX1 mRNA expression. Negative correlations were observed for cg00501904 (r = −0.556), cg03138928 (r = −0.526), cg05385434 (r = −0.435), and cg02453929 (r = −0.372), all statistically significant. These effect sizes are consistent with the interpretation that methylation at these loci contributes to transcriptional repression, although the cross-sectional design cannot establish causality. Nevertheless, the pattern is coherent with the broader literature on methylation-mediated silencing of detoxification and tumour-suppressor genes in HNSCC, including well-documented examples such as MGMT, in which promoter hypermethylation is associated with loss of protein expression and adverse clinical outcomes. (21–23) The present data suggest that EPHX1 may belong to this class of epigenetically silenced genes with prognostic relevance.

The survival analyses provide the most clinically translatable findings. High methylation at cg05935800 (log-rank P = 0.0016), cg02453929 (P = 0.044), and cg24868305 (P = 0.048) was associated with significantly worse overall survival. The cg05935800 result is especially notable because it is the only CpG in our dataset that satisfies three criteria simultaneously: tumour-specific hypermethylation, inverse correlation with expression, and significant association with survival. This convergence strengthens the case for cg05935800 as a candidate prognostic biomarker and suggests that epigenetic silencing of EPHX1 may identify a subset of HNSC patients with particularly aggressive disease. These findings align with accumulating evidence that DNA methylation signatures can stratify HNSCC patients by outcome, and they support the broader premise that methylation-based biomarkers may complement existing clinical and pathological prognostic factors. (14,15,24,25)

The clinical relevance of EPHX1 methylation is further underscored by the growing interest in epigenetic-modifying agents as therapeutic strategies in solid tumours. DNA methyltransferase (DNMT) inhibitors, such as azacitidine and decitabine, are approved for hematologic malignancies and are actively being investigated in head and neck squamous cell carcinoma (HNSCC), including combination strategies with immune checkpoint inhibitors and chemotherapy. Histone deacetylase (HDAC) inhibitors represent a complementary approach that can convert chromatin to a more open state and facilitate the re-expression of epigenetically silenced genes. Valproic acid (VPA), a well-established HDAC inhibitor, has demonstrated antineoplastic effects, and with combination therapies often showing superior results compared to monotherapy. Notably, in oral squamous cell carcinoma—a subtype of HNSCC—the combination of the DNMT inhibitor zebularine and the HDAC inhibitor VPA synergistically reduced tumour growth and reactivated tumour suppressor genes, accompanied by reduced DNA methylation in treated tumours. (26–28) These observations support the concept that reversing pathological methylation patterns, such as the tumour-specific EPHX1 hypermethylation at cg03138928 and cg05935800 identified in this study, may restore detoxification capacity and improve therapeutic responses. Future studies should explore whether EPHX1 methylation status could serve as a predictive biomarker for sensitivity to DNMT and HDAC inhibitors in HNSC, and whether combination epigenetic therapy can re-express EPHX1 and enhance the efficacy of conventional chemoradiation.

Several limitations warrant consideration. The analysis is observational and cross-sectional, precluding causal inference regarding the direction of the methylation–expression–survival relationships. The TCGA-HNSC cohort, while large and well-annotated, is predominantly composed of HPV-negative tumours and may not fully represent the changing epidemiologic landscape of HNSC. Tumour purity was not explicitly modelled in all analyses, and stromal contamination could influence methylation estimates. Smoking exposure was limited to self-reported years smoked, without detailed information on smoking intensity, cessation, or product type. Finally, the absence of functional validation means that the mechanistic basis for the observed associations remains inferential.

In summary, this study demonstrates that EPHX1 methylation in HNSC is a multi-dimensional phenotype shaped by tobacco exposure, tumour-specific epigenetic reprogramming, and transcriptional regulation. The identification of cg05935800 as a methylation marker associated with both reduced EPHX1 expression and poor survival provides a rationale for future functional studies examining whether EPHX1 silencing contributes causally to HNSC aggressiveness, and for validation of this CpG as a prognostic biomarker in independent cohorts.

## Conclusions

EPHX1 methylation is strongly associated with the transcriptional silencing of the gene in HNSC. Tumour-specific hypermethylation at loci such as cg03138928 and cg05935800 serves as a potential prognostic biomarker for poor overall survival, while smoking duration is associated with locus-specific hypomethylation in the tumour microenvironment. These findings highlight the EPHX1 epigenetic axis as a potential therapeutic target or biomarker in head and neck cancer.

## Acknowledgments

The results published here are in whole or based upon data generated by the TCGA Research Network: https://www.cancer.gov/tcga.

